# Sustained perturbation in functional connectivity induced by cold pain

**DOI:** 10.1101/633263

**Authors:** Elena Makovac, Ottavia Dipasquale, Jade B Jackson, Sonia Medina, Owen O’Daly, Jonathan O’Muircheartaigh, Alfonso de Lara Rubio, Steven CR Williams, Stephen B McMahon, Matthew A Howard

**Author notes:** Corresponding author: Dr Elena Makovac. King’s College London, Institute of Psychiatry, Psychology and Neuroscience, Department of Neuroimaging, Box 89, De Crespigny Park, London SE5 8AF, UK.

## Abstract

Functional connectivity (FC) perturbations have been reported in multiple chronic pain phenotypes, but the nature of reported changes is varied and inconsistent between cohorts. Increases and decreases in connectivity strength in task negative and positive networks, for example, the default mode and salience networks (DMN/SN), respectively, have been described, but how other networks are effected, for example, descending pain control networks, remains unknown. Whether connectivity changes relate to peripherally-mediated nociceptive afferent input, represent coping strategies or are sequelae of chronic pain, e.g. anxiety/depression, is also unknown. Here, we examined FC changes in response to experimentally-administered tonic cold pain in healthy volunteers as a means of disambiguating the nature of connectivity changes. We assessed FC prior to, during, and following tonic cold painful stimulation in four seed regions: ventromedial prefrontal cortex (vmPFC), rostral anterior insula (rAI), subgenual anterior cingulate cortex (ACC) and periaqueductal grey (PAG) and recorded subjectively reported pain using a computerised visual analogue scale. We saw DMN FC changes during painful stimulation and that inter-network communication between the rAI and sgACC seeds with the vmPFC became less anti-correlated during pain, whereas PAG-precuneus FC decreased. Pain-induced FC alterations largely persisted during a 6-minute recovery period following cessation of the painful stimulus. Observed FC changes related to the magnitude of individuals’ subjectively reported pain. We provide new insights into FC changes during and following tonic cold-pain and suggest that some FC changes observed in chronic pain patients may relate to the presence of an ongoing afferent peripheral drive.

## Introduction

Resting state functional magnetic resonance imaging (rs-fMRI) is a useful tool for investigating acute and chronic pain. Pain is an emergent property; a multi-system response to perceived threats to the body, comprising peripheral and central, autonomic, endocrine and immune components(Thacker and Moseley 2012). Accordingly, changes in functional connectivity (FC) from ‘task negative’, e.g. ‘default mode’ network (DMN) to ‘task positive’ networks, for example, the salience network (SN), including insula and anterior cingulate cortex (ACC)- have been reported in various chronic painful conditions (Baliki et al., 2014; (Schwedt et al., 2013; Seminowicz and Davis 2007a). Functional alterations in the periaqueductal grey (PAG), a crucial hub in descending pain control, have been reported in chronic pain, (Brooks and Tracey 2005; Hemington and Coulombe 2015), but reports of network perturbations vary between patient cohorts and are often inconsistent, for example, increased PAG-ventromedial PFC (vmPFC)/rostral ACC connectivity in low back pain (Yu et al., 2014), but reductions in the same regions in migraineurs(Jiang et al., 2016). Differences may simply reflect alternate clinical phenotypes, the influence of compensatory mechanisms, or sequelae such as depression or anxiety(Bair et al., 2008). Healthy volunteer studies facilitate examining pain mechanisms independently of these differences, but surprisingly, rs-fMRI studies evaluating tonic painful stimulation in these individuals remain relatively scarce. Reports of experimentally-induced tonic muscle pain(Alshelh et al., 2018) produced decreased oscillatory power in the main DMN hubs (posterior cingulate cortex-PCC, inferior parietal cortex, and vmPFC) and a pilot study of pain-inducing intramuscular hypertonic saline injection described insula-DMN connectivity changes (Zhang et al., 2014). To date, however, the effects of experimentally-induced noxious tonic stimulation in healthy volunteers on other pain-related functional networks, for example, endogenous pain control, or how networks interact with one another, remains to be assessed. Spontaneous, or tonic pain is a defining characteristic in chronic pain, but debate remains as to whether an ongoing peripheral afferent drive is necessary to produce it(Harte et al., 2017). FC changes observed in chronic pain may reflect the upstream effects of ongoing afferent peripheral nociceptive drive; in this study we investigated this using tonic cold pain in pain-free healthy volunteers.

Cold pain is an apposite choice. The cold pressor test is widely used to study pain and pain-associated autonomic reactions(Mourot et al., 2009). Cold allodynia is a frequent symptom in neuropathic pain(Bowsher and Haggett 2005) and abnormal cold perception is a major symptom in cold complex regional pain syndrome(Eberle et al., 2009). To date, only one evoked-response fMRI study has explored the neural basis of cold pain(La Cesa et al., 2014), reporting activations in ACC, thalamus, insula and PAG during repeated, short-duration cold-water hand immersion. How tonic cold pain modulates brain networks remains unknown. Here, we sought to determine changes in functional connectivity in pain-related networks in response to and following administration of a tonic, peripherally-mediated noxious cold stimulus. We hypothesised that cold pain would alter connectivity in vmPFC, rAI, PAG and sgACC functional networks and that the level of FC alteration would relate to individuals’ subjectively reported pain.

## METHODS AND MATERIALS

### Participants

Twenty healthy participants took part in the experiment (9 women, 11 men, mean age across the group = 26.05; SD = 5.32 years). Pain ratings were obtained from 18 participants only (two participants did not attend the final session). All participants were right-handed as assessed by the Edinburgh handedness inventory (Oldfield 1971). In addition to MRI contraindications, further exclusion criteria were: history of brain injuries, hypertension, neurological or psychiatric disease, and alcohol or drug abuse. To minimise the potential effects of menstrual cycle-related hormone fluctuations on pain responses (Vincent and Tracey 2010), female participants were all tested within the follicular phase of their menstrual cycle. Furthermore, to minimise the influence of diurnal variation on pain responses (Hodkinson et al., 2014; Strian et al., 2016) and on rs-fMRI network activity (Jiang et al., 2016), participants were always tested at approximately the same time for each visit. At the beginning of each visit, participants were tested for drugs of abuse (urine drug test) and alcohol consumption (alcohol breathalyser). All participants provided written informed consent. The study was approved by the National Research Ethics Service.

### Experimental procedure

Participants took part in an initial familiarisation session, followed by two identical scanning sessions and one post-scanning session. During the familiarisation session, participants became accustomed with the neuroimaging environment (in a mock scanner) and with the tonic cold-pain stimulation. The cold pain stimulation was delivered via an aluminium probe (4 × 20 cm), attached to the right inner forearm, through which cold water (2°C) was constantly circulated by means of two chillers. The constant circulation of the water ensured the stability of the temperature, which was also constantly monitored via a feedback loop. Participants were given a button-box to respond if the pain became intolerable.

During the scanning sessions (Session 2 and 3), participants underwent three 6-min resting state investigations: baseline (Pre-cold); cold-pain (Cold-pain) and post-cold recovery (Post-cold). During each resting state period, participants were instructed to rest with their eyes open, and keep their focus on the fixation cross presented at the centre of the screen, without thinking of anything and not falling asleep. To further explore the perception of cold pain, participants were tested during a post-scan session, where they were presented with the same 6-minute cold stimulation and a 6-minute post-cold interval. In this session, participants were asked to provide subjective ratings of the 6-minute cold and post-cold sessions on a visual analogue scale (VAS) ranging from 0 (indicating no pain) to 100 (indicating maximal imaginable pain in an experimental context).

### MRI acquisition and preprocessing

MR images were acquired on a GE MR750 scanner equipped with a 32-channel receive-only head coil (NovaMedical). Structural volumes were obtained using a high-resolution three-dimensional magnetization-prepared rapid gradient-echo sequence (TR = 7312 ms, TE = 3.02 ms, flip angle = 11⁰, slice thickness = 12 mm, 196 sagittal slices, FOV = 270 mm). Functional MRI data were collected with a T2*weighted multi-echo imaging (EPI) sequence sensitive to blood oxygenation level dependent (BOLD) signal (TR = 2 s, TE1 = 12 ms, TE2 = 28 ms; TE3= 44 ms; flip-angle 80°, 32 slices, 3mm slice thickness, 240 mm FOV, voxel size 3.75 × 3.75 × 3 mm).

Rs-fMRI datasets were pre-processed by combining different elements of established software toolboxes: AFNI (Cox 1996), the Advanced Normalization Tools (ANTs) (Avants et al., 2011) and FSL (Smith et al., 2004). By acquiring multiple echo images per slice, multi-echo fMRI permits identification of non-BOLD related sources of signal, preserving signals of interest (Dipasquale et al., 2017). This is of particular importance for pain studies, where pain-induced body movements or gross physiological changes may provide sources of artefactual non-BOLD-related signal alterations. Pre-processing steps were implemented in AFNI, and included volume re-alignment, time-series de-spiking and slice time correction. After pre-processing, functional data were optimally combined (OC) by taking a weighted summation of the three echoes using an exponential T2* weighting approach (Posse et al., 1999). The OC data were then de-noised with the Multi-Echo ICA approach implemented by the tool meica.py (Version v2.5 beta9, AFNI) (Kundu et al., 2013; Kundu et al., 2014), given its proven effectiveness in removing physiological and motion-related noise and increasing temporal SNR (Dipasquale et al., 2017; Kundu et al., 2013). Briefly, multi-echo principal component analysis was first used to reduce the data dimensionality in the OC dataset. Spatial ICA was then applied on one echo, and the independent component time-series were fitted to the pre-processed time-series from each of the three echoes to generate ICA weights for each echo. These weights were then fitted to the linear TE-dependence and TE-independence models to generate F-statistics and component-level κ and ρ values, which respectively indicate BOLD and non-BOLD weightings. The ρ metrics were then used to identify non-BOLD-like components to be regressed out of the OC dataset as noise. For further technical details on ME-ICA see (Kundu et al., 2015).

Using FSL, we regressed out the white matter (WM) and cerebrospinal fluid (CSF) signals, high-pass temporal filtered the data with a cut-off frequency of 0.005 Hz and spatially smoothed them with a 5 mm FWHM Gaussian kernel. Each participant’s dataset was co-registered to its corresponding structural scan with an affine registration and normalised to standard MNI152 space (with a non-linear approach) resampled to 2×2×2mm^3^ using ANTs.

#### Seed-Based fMRI Analysis

Anatomical ROIs were constructed using the Marsbar toolbox implemented in SPM 12 (http://marsbar.sourceforge.net/). We focused our interest in vmPFC as the central hub of the DMN, rAI as the central hub of the SN and often involved in chronic pain syndromes(Cottam et al., 2018), and the sub-genual ACC (sgACC) and PAG (both central hubs in the descending pain modulation pathway(Yu et al., 2014)). We created a 12 mm-radius sphere for the vmPFC (MNI 0 50 −8), 5 mm-radius sphere for the rAI (MNI= −36 30 4), 10 mm-radius sphere for the sgACC (MNI= 3 15 −10) and 3 mm-radius sphere for the PAG (MNI= 0 −30 −12). The volumes of the spheres differed according to the anatomical constraints imposed by each region. Coordinates were selected based upon those reported in the literature (Schweinhardt et al., 2006; Tzourio-Mazoyer et al., 2002; Yu et al., 2014). The average resting state fMRI time-series in each ROI was extracted for each participant and scan and used as a regressor in a 1^st^ level SPM analysis, to explore the network of areas associated with our seed regions.

Next, the first-level contrast images were entered into a flexible factorial design, with Condition (Pre-cold, Cold-Pain, post-cold) and Session (time 1, time 2) as main factors. Planned comparisons were performed for the following contrasts: Pre-cold vs Cold-pain, Pre-cold vs Post-cold and Cold-pain vs Post-cold.

#### Assessment of Relationships Between FC and Self-Reported Pain

To test for associations between FC in each seed region and subjective ratings of cold pain as indexed by VAS, a second level multiple linear regression general linear model (GLM) was computed. Inputs to the GLM were contrast images obtained at first level by subtracting cold-pain and pre-cold (ΔFC Cold-pain – Pre-cold) and between post-cold and pre-cold condition (ΔFC Post-cold – Pre-cold), with VAS ratings as covariates of interest. For illustrative purposes, parameter estimates for regions that showed significant correlations with VAS scores were extracted and plotted.

#### Statistical Inference

All mass-univariate voxelwise analyses were thresholded using an initial height threshold of Z = 2.58 and a Gaussian random field theory corrected (Worsley et al., 1992) cluster-significance threshold of p = 0.05.

## RESULTS

### Pain ratings

Following six-minute cold stimulation, participants gave an average pain rating of 44.94 (SD= 21.82) on a 0 – 100 VAS scale. At the end of the six-minute post-cold pain recovery phase, the participant pain intensity (rated on the VAS) was 1.50 (SD = 3.37).

### Functional connectivity networks

Seed-based FC of the vmPFC showed the core DMN network, consisting of the PCC/precuneus, bilateral parietal regions and PFC (Figure 1, A).

**Figure 1.**
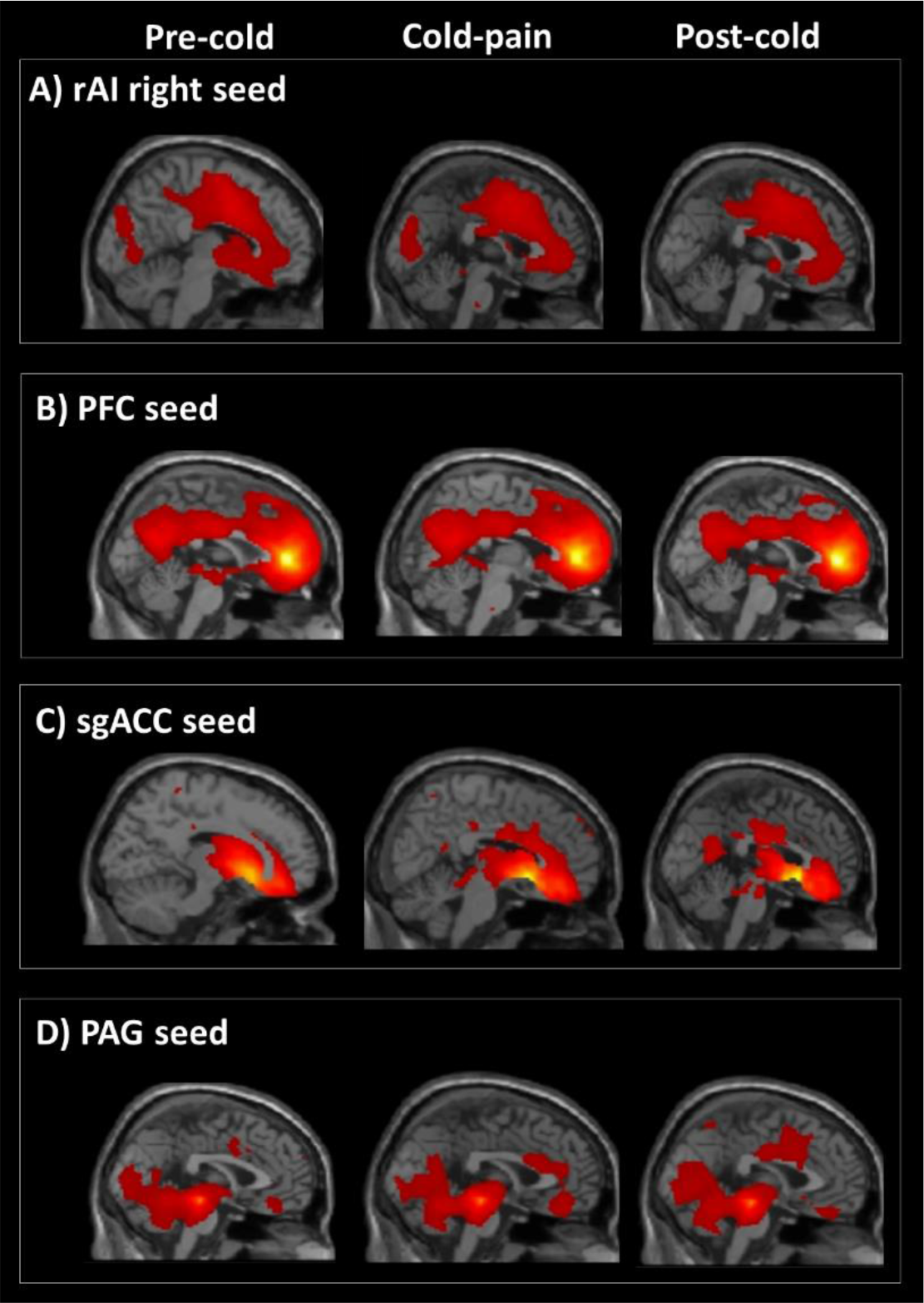
Resting-state FC networks for the four seed regions (vmPFC, rAI, sgACC, PAG). The networks are represented at rest (pre-cold), during cold pain and during post-cold pain recovery period. MNI [x y z] = 0 0 0.

Seed-based FC analysis of the rAI resulted in the identification of the SN network, comprising the following regions: sgACC and middle ACC, bilateral insula, bilateral supramarginal gyrus and cuneal cortex (Figure 1, B). Seed-based FC of the sgACC showed a network comprising the frontal medial cortex, thalamus, bilateral amygdala and bilateral parahippocampal gyrus (Figure 1, C). Seed-based FC of the PAG showed a network comprised of the bilateral parahippocampal/amygdalar areas, middle cingulate cortex, anterior insular cortex and prefrontal medial cortex (Figure 1, D), in line with other published studies[66].

### Functional connectivity of the vmPFC

When comparing Pre-cold to the Cold-pain condition, we observed an increase in FC between vmPFC and right dorsolateral prefrontal cortex (dlPFC) (Figure 2, A1). When comparing both Pre-cold and Cold-pain condition to Post-cold recovery condition, we observed a decrease in FC between vmPFC and the posterior areas of the DMN (lateral parietal lobules). When comparing Pre-cold to Post-cold only, we observed a decrease in FC between the vmPFC and precuneus/PCC and superior frontal gyrus, and an increase in FC between the vmPFC and lateral middle frontal gyrus/frontal pole and the ACC (Figure 2, A2 and A3; Table 1). In summary, the vmPFC increased its FC with other regions of the PFC (as part of the central executive network) and decreased its connectivity with more posterior regions of the DMN, and these alterations persisted in the post-cold phase.

**Figure 2.**
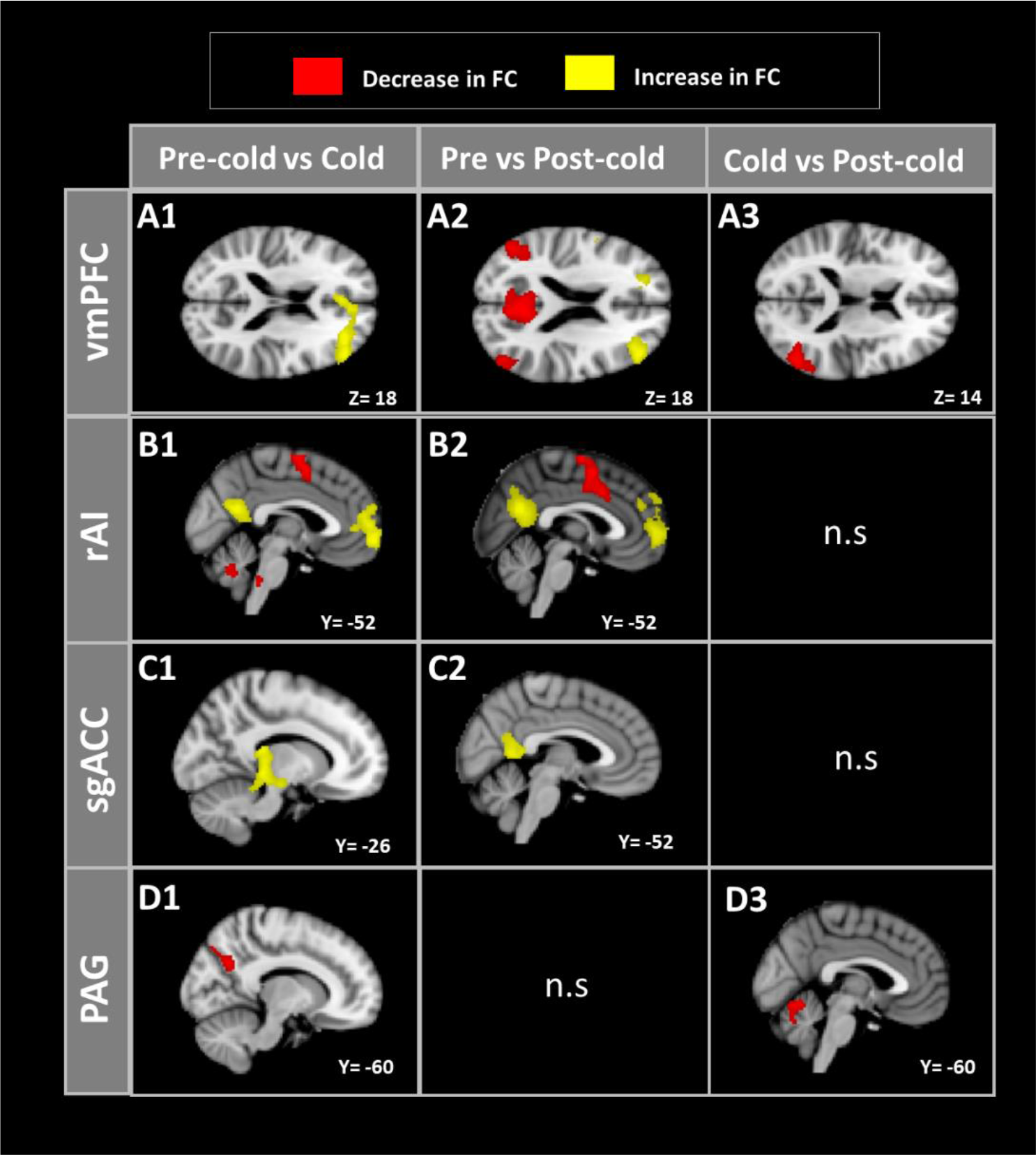
Functional connectivity alterations during cold pain and in the recovery phase. Graphical representation of brain areas whose FC was altered during cold pain and during post-cold pain recovery period. Yellow regions indicate an increase in FC between the seed regions (vmPFC, rAI, sgACC, PAG) and the rest of the brain; red areas indicate a decrease in the FC between the seed regions and the rest of the brain. In all four seeds (vmPFC, rAI, sgACC, PAG), the FC was altered when comparing cold-pain period to pain-free baseline (from top to bottom; A1-D1), and when comparing post-cold pain recovery period to baseline (from top to bottom; A2-D2). Less pronounced alterations were evident when comparing post-cold recovery period to the cold-pain period (from top to bottom; A3-C3), indicating that perturbation of the functional networks persisted during the 6 minutes following discontinuation of the painful cold stimulation. (n.s= non-significant results)

**Table 1.** Brain areas showing significant functional connectivity alterations during Cold-pain and Post-cold pain recovery period, when compared to Pre-cold baseline, for the vmPFC seed.

| Contrast | Brain area | Cluster |  | Voxel |  |
| --- | --- | --- | --- | --- | --- |
| | | $k$ | $p$ FWE | $T$ (F) | MNI $xyz$ |
| *Pre-cold < Cold-pain (T contrast)* |  |  |  |  |  |
|  | Dorso-lateral prefrontal cortex | 1442 | <0.001 | 4.27 | -30 42 20 |
| *Cold-pain > Post-cold (T contrast)* |  |  |  |  |  |
|  | Lateral occipital/parietal lobule | 733 | 0.016 | 4.47 | -50 -68 14 |
| *Pre-cold > Post-cold (T contrast)* |  |  |  |  |  |
|  | Precuneus | 3308 | <0.001 | 5.19 | -6 -58 18 |
|  | Lateral parietal lobule | 1187 | 0.001 | 4.57 | 44 -68 34 |
|  | Superior frontal gyrus | 657 | 0.027 | 4.09 | 16 28 50 |
| *Pre-cold < Post-cold (T contrast)* |  |  |  |  |  |
|  | Lateral middle frontal gyrus | 1211 | 0.001 | 4.52 | -44 40 20 |
|  | Lateral frontal pole | 2883 | <0.001 | 4.59 | 22 48 12 |
|  | Anterior cingulate cortex | 554 | 0.036 * | 5.00 | 12 48 12 |

\*Peak level FWE corrected

### Functional connectivity of the rAI

When comparing Pre-cold with Cold-pain, an increase in FC was observed between rAI and precuneus and vmPFC regions (both DMN hubs), and decreased FC between rAI, superior frontal gyrus and rostral-ventromedial medulla (RVM) (Figure 2, B1; Table 2). The same pattern was still observed when comparing Pre-cold with Post-cold (Figure 2, B2; Table 2); with an additional increase in FC with lateral parietal lobule, frontal orbital cortex and superior frontal gyrus. No differences in FC were observed when comparing Cold-pain with Post-cold pain. These data show that tonic noxious cold increases the FC between rAI and DMN areas and decreases the FC between rAI and the superior frontal gyrus and the descending pain control areas, and that this perturbation persists following the painful stimulation.

**Table 2.** Brain areas showing significant functional connectivity alterations during Cold-pain and Post-cold pain recovery period, when compared to Pre-cold baseline, for the rAI seed.

| Contrast | Brain area | Cluster |  | Voxel |  |
| --- | --- | --- | --- | --- | --- |
|  |  | *k* | *p FWE* | *T (F)* | *MNI xyz* |
| *Pre-cold < Cold-pain* |  |  |  |  |  |
|  | Precuneus/PCC | **835** | **0.006** | 4.34 | 8 -54 16 |
|  | vmPFC | **3006** | **<0.001** | 4.26 | 6 64 2 |
| *Pre-cold > Cold-pain* |  |  |  |  |  |
|  | Superior frontal gyrus | **950** | **0.003** | 4.27 | -26 2 64 |
|  | RVM | **1114** | **0.001** | 3.47 | 8 -40 -42 |
| *Pre-cold < Post-cold* |  |  |  |  |  |
|  | Precuneus/PCC | **2809** | **<0.001** | 5.65 | -4 -62 20 |
|  | Superior frontal gyrus | **1235** | **<0.001** | 5.05 | -22 22 36 |
|  | Lateral parietal lobule | **739** | **0.002** | 4.69 | -38 -72 34 |
|  | Frontal orbital cortex | **2129** | **<0.001** | 4.90 | 8 60 0 |
| *Pre-cold > Post-cold* |  |  |  |  |  |
|  | Premotor cortex | **1709** | **<0.001** | 4.70 | 4 4 56 |
|  | Cerebellum, left VI | **643** | **0.023** | 4.31 | -32 -58 -30 |

PCC= posterior cingulate cortex; vmPFC= ventromedial prefrontal cortex; RVM= rostral ventromedial medulla.

### Functional connectivity of the sgACC

When comparing Pre-cold with Cold-pain condition, we observed an increase in FC between sgACC and posterior thalamus (Figure 2, C1; Table 3, A). When comparing Pre-cold to Post-cold, we observed an increase in FC between sgACC and parahippocampal gyrus and the precuneus, and a decrease between the sgACC and a cluster comprising frontal orbital cortex and extending to the right anterior Insula and the putamen (Figure 2, C2; Table 3, A). No differences were evident when comparing Cold-pain condition to Post-cold recovery period.

**Table 3.**
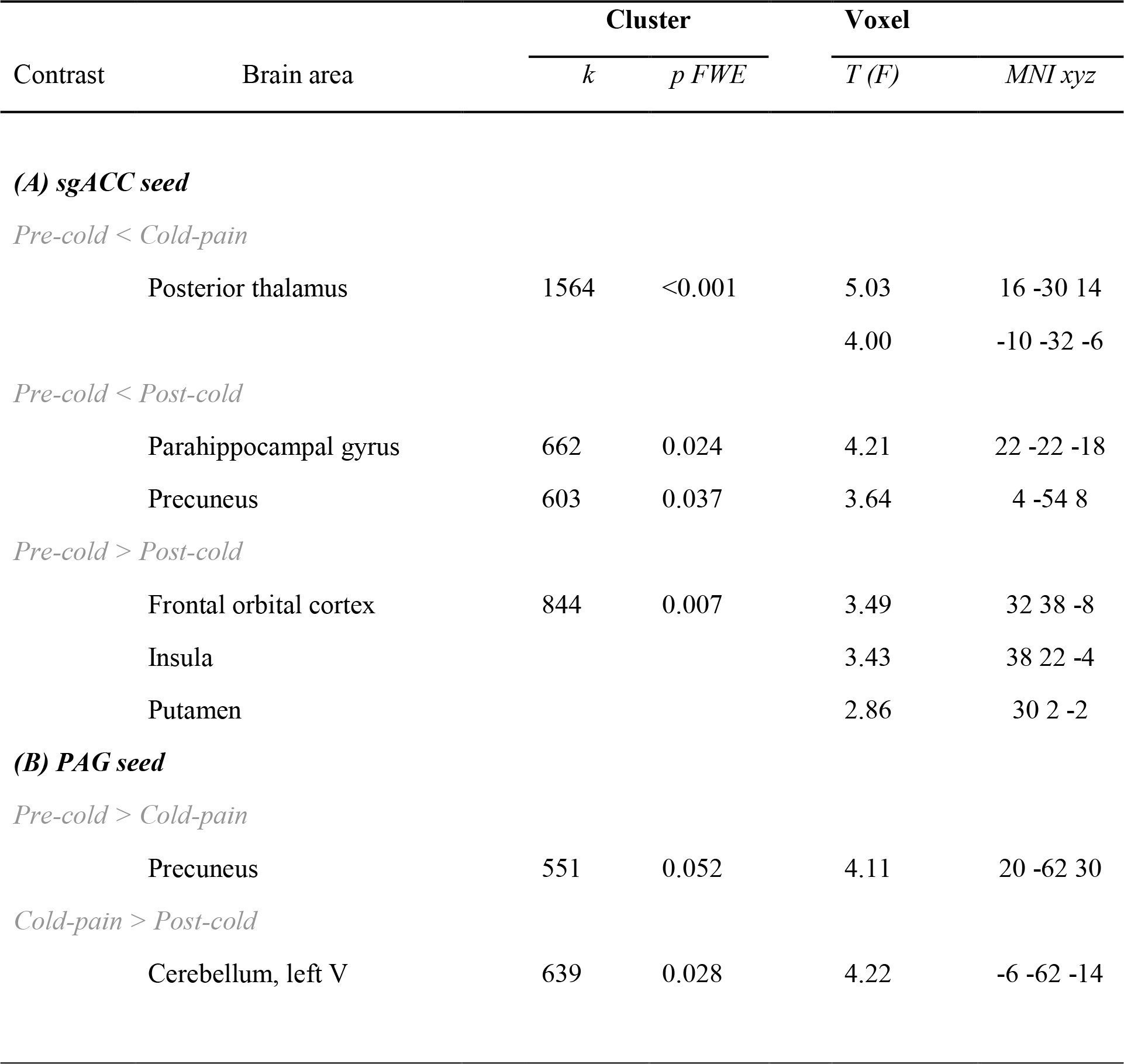
Brain areas showing significant functional connectivity alterations during Cold-pain and Post-cold pain recovery period, when compared to Pre-cold baseline, for the sgACC and PAG seed.

| Contrast | Brain area | Cluster |  | Voxel |  |
| --- | --- | --- | --- | --- | --- |
| | | $k$ | $p$ $FWE$ | $T$ ($F$) | $MNI$ $xyz$ |

|  |  |  |  |  |
| --- | --- | --- | --- | --- |
| Posterior thalamus | 1564 | <0.001 | 5.03 | 16 -30 14 |
|  |  |  | 4.00 | -10 -32 -6 |

|  |  |  |  |  |
| --- | --- | --- | --- | --- |
| Parahippocampal gyrus | 662 | 0.024 | 4.21 | 22 -22 -18 |
| Precuneus | 603 | 0.037 | 3.64 | 4 -54 8 |

|  |  |  |  |  |
| --- | --- | --- | --- | --- |
| Frontal orbital cortex | 844 | 0.007 | 3.49 | 32 38 -8 |
| Insula |  |  | 3.43 | 38 22 -4 |
| Putamen |  |  | 2.86 | 30 2 -2 |

|  |  |  |  |  |
| --- | --- | --- | --- | --- |
| Precuneus | 551 | 0.052 | 4.11 | 20 -62 30 |

|  |  |  |  |  |
| --- | --- | --- | --- | --- |
| Cerebellum, left V | 639 | 0.028 | 4.22 | -6 -62 -14 |

### Functional connectivity of the PAG

When comparing the Pre-cold condition with Cold-pain, an increase in FC was evident between the PAG and the precuneus (Figure 2, D1 and Table 3, B). When comparing Cold-pain to Post-cold, an increase in FC was evident between the PAG and an area of the cerebellum (left V; Figure 2, D3 and Table 3, B).

### Relationships Between FC and Self-Reported Pain

We conducted a whole-brain correlational analysis with VAS score as a regressor to investigate FC perturbations directly associated with the subjective perception of pain. We observed a negative correlation between cold-pain VAS ratings and the change in FC between the vmPFC and the precuneus/PCC area, when comparing both cold-pain and post-cold pain recovery phase with pre-pain condition (Figure 3, A), and a positive correlation was observed between the pain scores and the same contrasts (ΔFC Cold-pain-Pre-cold and ΔFC Post-cold – Pre-cold), for the FC between PFC and dorsolateral PFC (Figure 3, Table 4). These results indicate that individuals reporting a higher subjective perception of cold pain showed a greater functional decoupling between prefrontal and posterior areas of the DMN and a stronger increase in FC between vmPFC and lateral areas of the PFC.

**Figure 3.**
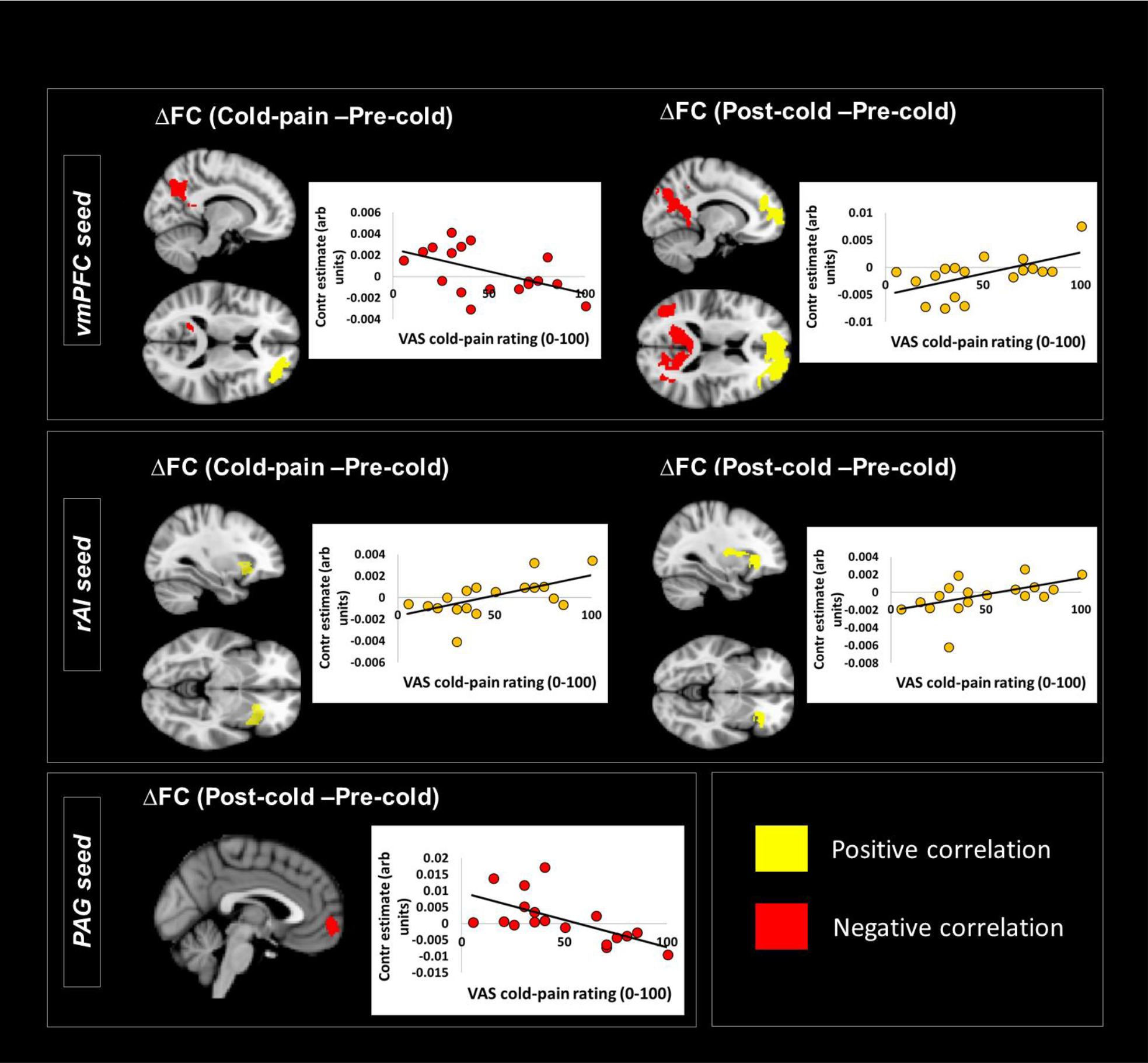
Correlation between changes in functional connectivity and pain ratings. Brain areas showing a significant association between the 6-minutes cold-pain VAS rating and the change in functional connectivity between Pre-cold and Cold-pain condition (ΔFC Cold-pain – Pre-cold) and pre-cold and post-cold (ΔFC Post-cold – Pre-cold), for the three seed regions of interest: PFC (A), rAI (B) and PAG (C).

**Table 4.**
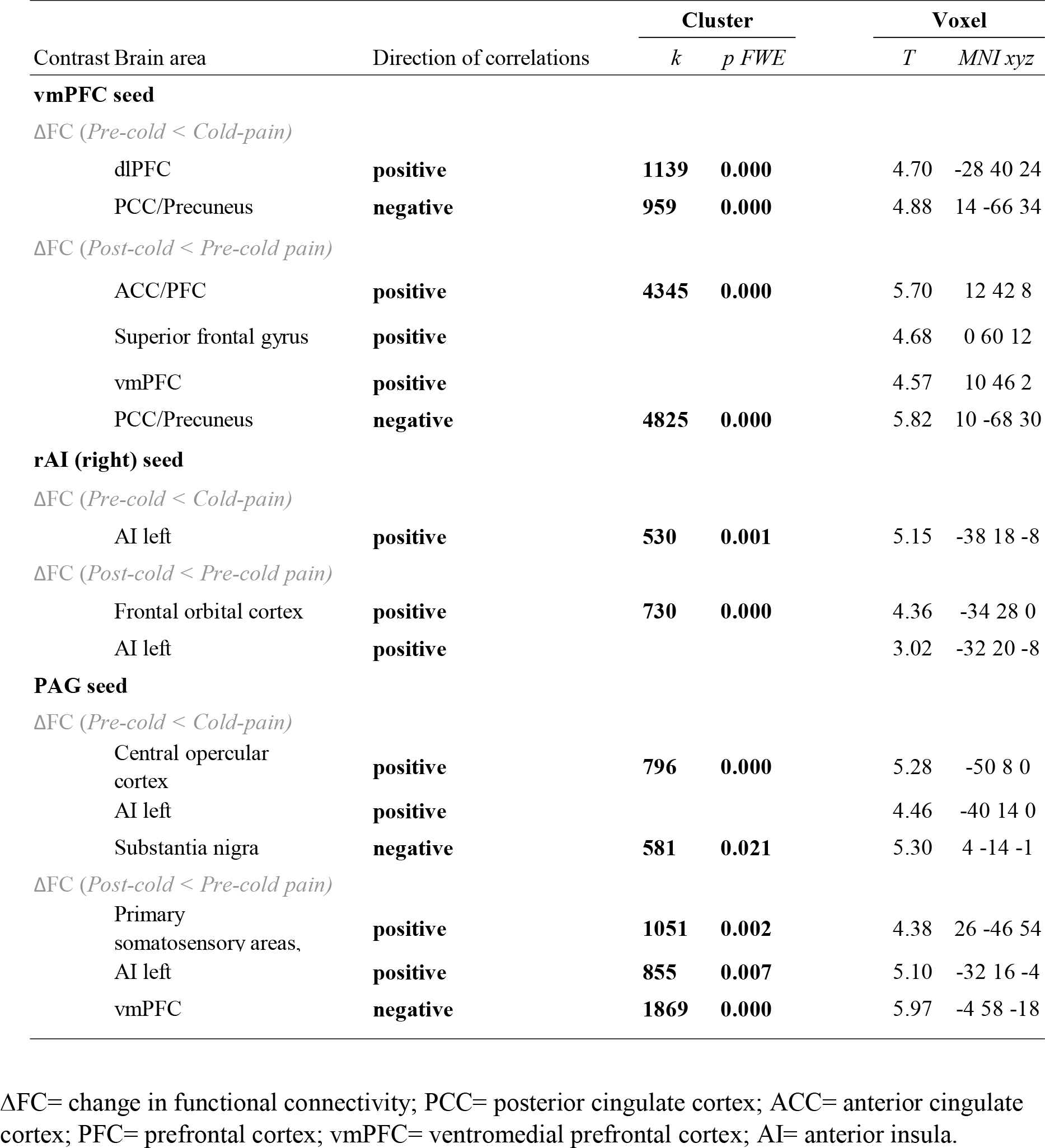
Brain areas showing significant functional connectivity associations with subjective pain ratings during the 6 minutes cold painful stimulation.

A positive correlation was observed between pain ratings and the FC between the right rAI and the left rAI, when comparing both the Cold-pain and Post-cold pain condition with the Pre-cold (Figure3, B, Table 4). This result indicates that individuals whose insular areas exhibited the strongest increase in FC during cold stimulation experienced the highest pain.

A positive correlation was observed between pain ratings and the change in FC (Cold-pain vs Pre-cold) between the PAG and the frontal opercular/anterior insular cortex, indicating that an increased functional coupling between these areas is associated with lower levels of experienced pain (Figure 3, C; Table 4).

A negative correlation was observed between pain ratings and the change in FC (Cold-pain vs Pre-cold) between the PAG and the SN/VTA, indicating that strongest interaction between these areas occurred when lowest levels of pain were reported. Further, subjective pain ratings correlated positively with the change in FC during the recovery phase (Post-cold vs Pre-cold) between the PAG and the left anterior insula and temporal cortex and right primary somatosensory area. Negative correlations with the change in FC (Post-cold vs Pre-cold) between the PAG and the vmPFC indicated that higher experienced pain was associated with a reduction in functional communication between PAG and prefrontal areas and an increase between PAG and insula and somatosensory areas (Figure 3, C; Table 4).

## DISCUSSION

Rs-fMRI has shown great potential in the investigation of chronic-pain related FC alterations(Yoshino et al., 2017). However, the degree to which these reports of changes in transient, state-dependent interactions between brain regions reflect the chronic pain experience per se, or more generalised network responses to highly salient, tonic noxious stimulation remains uncertain. In this healthy volunteer study, we used a tonic 6-minute noxious cold stimulus with the aim of inducing FC perturbations in the main pain hubs (vmPFC, rAI, PAG and sgACC). The effects of ongoing cold pain elicited alterations in the default mode, salience, central executive and descending pain control networks, several of which persisted for several minutes following stimulation. Importantly, pain-induced network alterations were associated with participants’ subjectively reported ratings of their ongoing pain experiences. We interpret these cold-induced network changes as responses by stimulus salience, interoceptive, autonomic and endogenous pain control systems to homeostatic threat and demonstrate that FC alterations previously observed in patients with chronic pain can be induced by ongoing tonic peripheral noxious stimulation in healthy individuals.

We demonstrate that tonic cold painful stimulation increased the FC between major hubs of the central executive network (CEN), the vmPFC and dlPFC(Chen et al., 2013). Further, during the Post-cold recovery phase, FC between vmPFC and posterior areas of the DMN, the precuneus/PCC and lateral parietal lobules, decreased. These findings echo reports of decreases in FC between the PFC and precuneus in several different chronic pain conditions, including chronic back pain, complex regional pain syndrome and knee osteoarthritis(Baliki et al., 2014). Similarly, increased FC between the dlPFC and the bilateral rostral ACC and vmPFC has also been reported in fibromyalgia(Kong et al., 2018). In healthy individuals, the dlPFC has a causal regulatory influence on the vmPFC(Chen et al., 2013), where an engagement of the CEN by an active task temporarily ‘shuts-down’ the DMN. Our data demonstrate that painful stimulation disrupts this inhibitory control from the CEN, increasing the FC between these two networks. This inability to down-regulate the DMN by the CEN when an individual is in pain might (at least in part) explain decision-making and executive disabilities in chronic pain patients(Baker et al., 2016).

Connectivity decreases between vmPFC and PCC and increases between vmPFC and dlPFC were associated with cold-pain ratings. Increased PAG-vmPFC connectivity in the post-cold period correlated negatively with pain ratings, indicating that individuals with the highest experienced pain exhibited lower FC among these two areas during the recovery. A similar relationship was previously described in patients with chronic low back pain(Yu et al., 2014). These findings engender the working hypothesis that these patterns of FC in chronic-pain populations may be due to the presence of peripherally mediated afferent signalling, rather than being associated with other chronic pain sequelae such as anxiety, depression or sleep deprivation, or utilisation of alternative coping mechanisms(Simons et al., 2014).

We observed a decrease in anti-correlation between the rAI, a core node in the salience network, and the main hubs of the DMN during cold painful stimulation; these relationships persisted in post-pain recovery phase. Increased SN – DMN connectivity is also a result widely reported within the chronic pain literature. For example, increased connectivity between the DMN and the insula has been described in patients with diabetic neuropathic pain(Chen et al., 2013) and patients with rheumatoid arthritis(Flodin et al., 2016). Increased insula-DMN coupling decreases after treatment in fibromyalgia patients and is associated with a reduction in pain(Napadow et al., 2010). Our data, together with the rest of the chronic-pain literature, support a critical role of the rAI in switching between the DMN and the CEN during the elaboration of salient stimuli including pain(Sridharan et al., 2008). During cold pain, the rAI exhibited decreased connectivity with the RVM. The rAI and the RVM are both involved in the descending pain modulatory pathway, and their reciprocal anatomical connections mediate autonomic reactions(Kapp et al., 1985). Our results suggest an engagement of descending pain control mechanisms in response to cold pain with the aim of re-establishing homeostasis.

Both the sgACC and the PAG showed connectivity changes with the precuneus/PCC area during and following noxious cold stimulation, suggesting that the PCC/precuneus region acts as a hub for the interaction among functional networks during pain. The PCC/precuneus area is engaged during the assessment of the emotional relevance of stimuli(Vogt 2005) and its activity varies with arousal states(Vogt and Laureys 2005). Individuals with the largest increase in precuneus response during pain have lower pain sensitivity(Goffaux et al., 2014) and pain perceived during the application of non-noxious stimuli (i.e. allodynia) is associated with increased activity in the precuneus(Witting et al., 2001). Individuals with traumatic neuropathic pain have also been reported to exhibit differential regional cerebral blood flow in the PCC(Hsieh et al., 1995). A failure to down-regulate the PCC (and the associated DMN) is related to cognitive dysfunctions, both in healthy subjects and clinical populations(Weissman et al., 2006). We suggest that our observations of decreased rAI-PCC anticorrelation and increased sgACC – Precuneus connectivity indicate that pain interferes with DMN control, and that these perturbations contribute to the complex and intimate relationships between pain and cognition(Seminowicz and Davis 2007b).

Cold pain ratings correlated negatively with the change in PAG-SN/VTA FC during pain. Both the VTA and SN are part of the mesolimbic dopamine system, implicated in motivated behaviour(Taylor et al., 2016). The role of the VTA in pain is also well-characterised. In animals, stimulation of this area increases pain thresholds and decreases spinal dorsal horn activity(Li et al., 2016). In humans, VTA dopaminergic neurons are thought to be responsible for the dopamine release after aversive stimuli, such as psychosocial stress or pain(Taylor et al., 2016). The ventrolateral PAG contains dopaminergic neurons which are thought to underlie opioid-mediated antinociception. Therefore, the functional association between the PAG and the SN/VTA might indicate an involvement of dopamine-mediated descending pain inhibition(Bannister and Dickenson 2016), where individuals with the strongest increase in connectivity between these areas reported lower perception of pain. Our findings support also the role of this area in reward and pain relief(Navratilova and Porreca 2014).

Subjective pain ratings were also positively associated with increased FC between the right and left rAI and the PAG during cold pain. Although the insula is often bilaterally activated during noxious somatosensory stimulation(Coghill et al., 1999), functional lateralization has also been observed in relation to autonomic reactions, where direct stimulation of the right and left insula produces sympathetic and parasympathetic reactions, respectively(Oppenheimer et al., 1992). We interpret that increases in connectivity between PAG/right rAI and left rAI might indicate an attempt to recover homeostasis via activation of the parasympathetic nervous system. rAI is also a critical structure involved in *interoception*, the ability to accurately perceive signals from the body. Interoceptive accuracy mediates pain perception and is impaired in chronic pain conditions(Di Lernia et al., 2016). We speculate that increased FC between PAG/right rAI and left rAI may underpin linking between meta-cognitive and emotional aspects of pain (via the right rAI) with autonomic and endogenous pain modulating functions (via the PAG and left rAI). Reciprocal projections between rAI and the PAG (via the amygdala and thalamic nuclei) support this theory, providing circuitry that during pain may elicit emotional arousal and trigger the descending pain modulatory system(Moraga-Amaro and Stehberg 2012).

We observed that cold-induced rs-fMRI perturbations persisted in several functional networks during the post-cold condition. These findings accord with reports of acute psychosocial stressors eliciting prolonged, post-stressor DMN connectivity responses(Vaisvaser et al., 2013). and studies reporting BOLD responses that do not immediately return to baseline following the completion of both cognitively demanding tasks(Barnes et al., 2009), and simpler paradigms such as pressing a button for a fixed duration(Tung et al., 2013). In the DMN and PAG networks, post-cold perturbations were directly associated with pain ratings, indicating that the persistence of FC alterations is not a generalised rs-fMRI phenomenon. We urge caution to other pain neuroimaging researchers that the findings of ‘back-to-back’ fMRI assessments featuring tonic painful stimuli may well be confounded.

In summary, we have provided new insights regarding the neural representation of tonic cold-pain and demonstrate that connectivity alterations observed in chronic pain patients, (notably, those involving the DMN, SN and areas functionally associated to the PAG and sgACC), can be elicited by tonic, noxious, peripheral afferent stimulation in healthy volunteers. Our data may guide future interpretations of chronic-pain related rs-fMRI alterations, helping disentangle pain-related alterations from functional abnormalities associated with chronic pain comorbidities. We tentatively suggest our results may also offer input to *nociplastic* definitions of chronic pain(Kosek et al., 2016); rs-fMRI network descriptions may facilitate future disambiguation of peripherally-mediated, from central nervous system maintained, chronic pain states. Identifying aetiological pain network ‘neurosignatures’ offers the enticing potential promised by precision medicine that identify specific pathological pain mechanisms operating in individual patients(Rosa and Seymour 2014). Our findings may in the long term help the direction of focussed treatment approaches and inform the development of new and much-needed therapies.

## Acknowledgments

This work was funded by a Medical Research Council Experimental Medicine Challenge Grant (MR/N026969/1). MAH, SM and SW are also supported by the NIHR Biomedical Research Centre for Mental Health at the South London and Maudsley NHS Trust. JOM is supported by a Sir Henry Dale Fellowship jointly funded by the Welcome Trust and the Royal Society (Grant Number 206675/Z/17/Z) and a Medical Research Council (MRC) Centre grant (MR/N026063/1). We thank Simon Hill for both the development and calibration of the stimulation equipment. Declaration of interest: None.

